# Speech clarity shapes auditory attention and visual-signal coupling during multimodal sentence comprehension

**DOI:** 10.64898/2026.07.13.738151

**Authors:** Cecília Hustá, Noor Seijdel, Linda Drijvers

**Affiliations:** Max Planck Institute for Psycholinguistics, Nijmegen, The Netherlands; Donders Institute for Brain, Cognition and Behaviour, Radboud University, Nijmegen, The Netherlands

**Keywords:** multimodal language, gesture, informativeness, speech clarity, integration, RIFT

## Abstract

Face-to-face communication requires listeners to attend, integrate, and weigh multiple communicative signals, including auditory speech, mouth movements, and co-speech gestures. The contribution of these signals may depend on the reliability of auditory input and the informativeness of the available signals. We utilized rapid invisible frequency tagging (RIFT) with EEG to examine how participants attend to and integrate these different signals in clear and adverse listening conditions. Participants watched videos of an actress producing clear or noise-vocoded sentences. Auditory speech was amplitude-modulated at 58Hz, while the luminance of the gesture and mouth regions was frequency-tagged at 63Hz and 65Hz. Degraded speech elicited stronger responses at the auditory tagged frequency, suggesting increased attentional gain to the auditory signal when listening was challenging. In contrast, clear speech elicited stronger responses at the gesture tagged frequency and a stronger 2Hz intermodulation response (65-63Hz), reflecting enhanced nonlinear coupling between mouth movements and gestures. Finally, in degraded speech, the informativeness of mouth movement, but not gesture, was associated with intermodulation strength, suggesting that the informativeness of mouth movements plays a greater role in multisensory interaction when listening is challenging. Our findings demonstrate that both signal reliability and informativeness shape multisensory integration during spoken language comprehension.

## Introduction

Listeners are remarkable at extracting information from naturalistic communication, integrating auditory speech signals with visual cues such as mouth movements and gestures to construct meaning and support successful comprehension. The extent to which listeners rely on these different sources of information depends on both environmental conditions and the informativeness of the available signals, defined as the extent to which a single-modality signal provides information about the intended linguistic message. For example, in a noisy café, a listener may rely heavily on a speaker’s mouth movements and gestures to compensate for degraded auditory input. Even when speech is clear, gestures can contribute additional semantic information that is not conveyed in speech (e.g., making loose fists and moving them in a rhythmic circular motion while uttering the word *went*, as if driving). Although previous research has demonstrated that both mouth movements and gestures facilitate speech understanding (e.g., Drijvers & Özyürek, 2017), less is known about how listeners allocate attention across these signals when their relative informativeness varies (but see Krason et al., 2022; Zhang et al., 2021). In the present study, we investigated how auditory degradation and the relative informativeness of gestures and mouth movements influence attentional allocation to these signals and their interaction during speech comprehension.

In addition to listening, interlocutors process visual signals generated by the face and body, particularly in face-to-face communication. Listeners naturally attend to speaker’s face, making information conveyed through mouth movements, facial expressions, head movements readily available during language comprehension (Argyle et al., 1994; Gullberg & Holmqvist, 1999). But, visual signals are not limited to the face: a large-scale analysis of an English TV news archive found that gestures accompanied approximately 62% of utterances (Wilding et al., 2025). Critically, attention to gestures is pervasive, with participants attending to the gestures even when they are incongruent with speech (Arnheim & McNeill, 1994; Kelly et al., 2010), albeit often through the periphery of their vision. During clear speech comprehension, listeners tend to fixate on the face of a speaker for about 90-95% of the time (Argyle & Cook, 1976; Argyle & Graham, 1976; Gullberg & Kita, 2009), and tend not to fixate on gestures directly (Gullberg & Holmqvist, 2006), implying that they can extract semantic information from them without directly gazing at them. Similarly, during degraded speech comprehension, overt visual attention is often directed at the mouth, and not at gestural information, while covertly this information is being attended to (Drijvers, Vaitonyte, Ozyurek, 2019; Seijdel et al., 2024). In general, research suggests that the presence of visual signals improves speech perception, with performance often being highest when multiple signals (e.g., mouth movements and gestures) are available (Drijvers & Özyürek, 2017).

Visual signals can support comprehension on different levels. On a lexical level, seeing a speaker’s mouth movements facilitates speech recognition (Peelle & Sommers, 2015) by aiding phonological processing (Mcgurk & Macdonald, 1976; Sumby & Pollack, 1954) or by reducing lexical competition (Jesse & Massaro, 2010; Lachs & Pisoni, 2004). On a semantic level, gestures, especially iconic gestures, facilitate comprehension (Dargue et al., 2019; Hostetter, 2011; Wu & Coulson, 2015), possibly by aiding semantic processing (McNeill, 1992). This aligns with the neuroimaging findings of Skipper and colleagues (2009) who showed that mouth movements and co-speech gestures facilitate language comprehension through partly distinct neural pathways: mouth movements engage motor and phonological processing networks involved in speech perception, whereas gestures engage motor and semantic processing networks involved in meaning extraction. Interestingly, facilitation of visual signals seems more pronounced under adverse listening conditions (for mouth movements: Ma et al., 2009; Ross et al., 2006; Schwartz et al., 2004; Sumby & Pollack, 1954; for gestures: Drijvers & Özyürek, 2017; Holle et al., 2010; Obermeier et al., 2012). Together, these findings demonstrate that visual signals support speech comprehension and this interaction is modulated by the reliability of the auditory input.

Listeners are unlikely to rely on the various signals in a fixed or uniform way; instead, the contribution of each signal depends on various contextual factors. For example, Krason and colleagues (2022) found that audiovisual word recognition was shaped by the informativeness of visual signals. Highly informative congruent gestures facilitated recognition, whereas highly informative incongruent gestures produced the strongest interference. Importantly, informative mouth movements mainly aided recognition under noisy conditions. Zhang and colleagues (2021) demonstrated that visual signals influence EEG markers of semantic processing, with effects arising from dynamic interactions among signals rather than from independent, additive processing. Together, these findings indicate that audiovisual speech comprehension is not only shaped by the informativeness of individual signals, but also by dynamic interactions that determine how these signals are weighted over time. If listeners prioritize the most informative signals during comprehension, then differences in signal informativeness should be reflected in the amount of attentional gain allocated to those signals, and/or in the strength of the interactions between the signals. Moreover, flexible signal weighting could be expressed in at least two separable ways. Listeners may increase attentional gain to an individual signal, such as just to speech, just to the mouth movements, or the gestures. Alternatively, they may change the degree to which different signals are coordinated or integrated. Distinguishing between these possibilities thus requires a method that can track responses to individual communicative signals, while also indexing interactions between those signals.

To investigate covert attentional allocation and stimulus interactions, recent studies have employed rapid invisible frequency tagging (RIFT; for review see Arora et al., 2025). This technique involves periodic modulation of visual luminance and auditory amplitude at frequencies above 50 Hz, generating reliable steady state evoked potentials (SSEPs). The amplitude of these responses serves as a neural marker of attention to the tagged visual and auditory stimuli (Toffanin et al., 2009). Previous studies have already shown that speech clarity influences attention to auditory and visual information. Auditory SSEPs were stronger when speech was degraded, indicating increased attention to the auditory signal under difficult listening conditions (Drijvers et al., 2021; Seijdel et al., 2024). Unlike auditory SSEPs, visual SSEPs elicited by gestures showed a more complex pattern. There are studies showing stronger SSEPs in clear speech (Drijvers et al., 2021) and sometimes in degraded speech (Seijdel et al., 2024), possibly depending on the differences in task demands or attentional allocation between the various signals. It is therefore crucial to investigate how attention towards mouth movements influences attentional allocation towards these signals.

Tagging two stimuli at different frequencies produces intermodulation (IM) frequencies, which arise from nonlinear interactions between the tagged signals (f₂ ± f₁; e.g., Regan et al., 1995), providing a neural measure of the interaction between the representations of the two stimuli (Arora et al., 2025; Seijdel et al., 2023). Previous studies have used the IM frequency to tap into interaction between audio-visual representations (Drijvers et al., 2021; Hustá et al., 2025; Seijdel et al., 2024). IM power was strongest when speech was clear and the conditions for gesture speech interaction were optimal (Drijvers et al., 2021). More recent findings showed a different pattern, namely, a stronger power at the IM frequency for degraded speech with matching gestures (Seijdel et al., 2024), depending on attentional allocation. Seijdel’s results suggest that IM responses may reflect not only lower-level interactions between gestures and speech but also the effects of higher level factors on the interaction dynamics.

In the present study, we employed RIFT to investigate how listeners integrate and attend to speech, mouth movements, and gestures, and how the informativeness of the visual signals influences the interactions between the different signals. Participants watched videos where an actress produced sentences containing critical verbs that were accompanied by gestures (Wilms et al., 2022). RIFT was utilized to modulate the amplitude of the speech at 58 Hz, while the luminance was modulated at 63 Hz in the area containing gestures and at 65 Hz in the area containing mouth movements. The speech was either noise-vocoded or clear. Gesture and mouth-movement informativeness was established in a pretest by determining how each of the visual signal independently predicted the target word (i.e., lexical affiliate).

Based on Drijvers and colleagues (2021), we expected higher auditory SSEPs at 58 Hz in the degraded compared to clear condition, and higher visual SSEPs at 63 Hz elicited by gestures in the clear compared to degraded condition. Considering that mouth movements and gestures support comprehension through different mechanisms (phonological vs., semantic level), we hypothesized that listeners would allocate more attention to mouth movements when speech was ambiguous, resulting in higher visual SSEPs at 65 Hz in the degraded compared to clear condition. We expected to observe IM frequencies resulting from the nonlinear interaction of the base tagging frequencies at 2 Hz, reflecting interactions between representations of mouth movements and gesture; at 5 Hz, reflecting interactions between representations of gesture and speech; and at 7 Hz, reflecting interactions between representations of mouth movements and speech. Based on Drijvers and colleagues (2021), we expected higher IM power when speech was clear rather than degraded, reflecting more efficient (two-way) interactions between the representations on speech, mouth movements, and gesture. Furthermore, we expected that higher gesture and mouth-movement informativeness would facilitate interaction between the respective visual signal and speech representations, and between the two visual signal representations.

## Methods

### Participants

Thirty-one native Dutch-speaking adults (Mean age = 24.76, SD = 4.39, 24 women) took part in the EEG experiment. One participant was excluded because they reported an ADHD diagnosis after completing the experiment, and five participants were excluded because of technical issues (missing data, EEG/RIFT issues), resulting in 25 participants in the final dataset. All included participants reported normal hearing, normal or corrected-to-normal vision, and no history of neurological, psychiatric, hearing, or developmental language disorders. Participants were recruited via the participant database of the Max Planck Institute for Psycholinguistics in Nijmegen and received monetary compensation for their participation. All participants gave written informed consent before the start of the experiment, and the study was approved by the local ethics committee.

### Stimuli and design

Participants were presented with short audiovisual sentence videos (n=170) from Wilms et al. (2022). In each video of about 2 seconds, an actress produced a Dutch sentence of approximately four to six words. Each sentence had a similar structure, with a finite verb in second position, and ending with an infinite action verb (e.g., ‘Zij zit vaak te huilen’, [she is often crying]). The action verbs were accompanied by an iconic gesture. For more details on the stimuli, and the corresponding pretests, see Wilms et al. (2022).

The speech in the videos were either clear or noise-vocoded (10-band noise-vocoding, see for pretests: Wilms et al., 2022). To achieve this, we first separated the audio files from the videos, sampled them at 44.1 k Hz, intensity-scaled them to 70 dB, denoised them, noise-vocoded them using ten logarithmically spaced frequency bands between 50 and 8000 Hz, and then recombined with the original videos, using *Praat* (Boersma & Weenink, 2022), following the same procedure as in previous studies (Drijvers & Ozyuek, 2017; Krason et al., 2022).

### Informativeness pretests

Gesture and mouth-movement informativeness were obtained from separate online pretests that were implemented in Gorilla (Anwyl-Irvine et al., 2020). All participants were native speakers of Dutch, reported normal hearing and no history of language, neurological or other disorders. None of the participants took part in the main experiment.

For the gesture informativeness pretest, 11 participants looked at videos that were cropped around the gesture region. On each trial, participants saw the gesture twice and were asked to type the verb that they thought best matched the hand movement that they saw. Gesture informativeness was quantified as the semantic similarity between each typed response and target word using Dutch fastText Common Crawl/Wikipedia word vectors (see Bojanowski et al., 2016; Grave et al., 2018). For each response-target pair, we calculated cosine similarity between the corresponding word vectors (similar to Kenter & de Rijke, 2015; van Paridon & Thompson, 2021). Response-level cosine similarities were then averaged per stimulus, resulting in one item-level gesture informativeness score.

For the mouth-movement informativeness pretest, 9 participants looked at videos that were cropped to around the face region. All videos were temporally trimmed around the target verb. Participants saw the video twice, and were asked to type the verb they thought the actress had produced. Mouth-movement informativeness was quantified as an item-level average of the orthographic distance as an approximation for phonologic distance between the response and the target, using the token-sort-ratio (Bosker, 2021).

Across the final item set, gesture informativeness and mouth-movement informativeness varied substantially across items (gesture: M = 0.49, SD = 0.24, range = 0.16–1.00; mouth movements: M = 46.08, SD = 10.55, range = 23.67–88.89).

### Procedure

Participants were tested individually in a dimly lit room and sat approximately 90 cm from the projection screen. The experiment was presented using MATLAB 2020b (Mathworks inc, Natrick, USA) and Psychophysics Toolbox (Brainard, 1997; Pelli, 1997; Kleiner et al., 2007). All stimuli were projected with a PROPixx DLP LED projector (VPIXX Technologies Inc, Saint-Bruno-de-Montarville, Canada) and using a GeForce GTX960 2GB graphics card with a refresh rate of 120 Hz. The PROPixx projector can achieve a presentation rate up to 1440 Hz by interpreting the four quadrants and three color channels of the GPU screen buffer as individual, smaller, grayscale frames, which are then presented in rapid succession, leading to an increase of a factor 12 (4 quadrants x 2 color channels x 120 Hz, User Manual for ProPixx, Vpixx Technologies Inc., Saint-Bruno-de-Montarville. Canada; see for more details and background: Arora, Husta et al., 2026).

Each trial, participants were first presented with a fixation cross for 1000 ms, the stimulus video clip (around 2000 ms), a 1500 ms delay, and then an answering screen where they had to type the sentence that was communicated in the video clip (‘Typ de zin hier: …’, [Type the sentence here: …]). Each participant saw each item once, either in clear or degraded speech, using semi-randomized stimulus lists. The entire experiment took about 1.5 hours, including preparation time.

### Frequency tagging

RIFT was used to tag the auditory speech signal, the gesture region, and the mouth region, following previous RIFT work that demonstrated that high-frequency tagging of auditory and visual signals can be used to track sensory responses and nonlinear IM responses during naturalistic settings (Drijvers et al., 2021; Seijdel et al., 2023; 2024; Husta et al, 2025). We modulated the amplitude of the speech signal with a 58 Hz sinusoid, and a modulation depth of 100%. We multiplied the luminance of the pixels in the gesture space and mouth movement space with a 63 Hz and a 65 Hz sinusoid (modulation depth = 100%, modulation signal equal to 0.5 at sine wave zero-crossing to preserve the mean luminance of the video). The three tagging frequencies thus yielded three different IM responses: 2 Hz for mouth movement-gestures coupling (65-63 Hz), 5 Hz for auditory speech-gesture coupling (63-58 Hz), and 7 Hz for auditory speech-mouth movement coupling (65-58 Hz).

### EEG acquisition and preprocessing

EEG was recorded using 32 electrodes (actiCap, Brain Products, Germany), arranged according to the 10-20 system. 30 electrodes were placed on the scalp, two electrodes were placed on the left and right mastoid and used for rereferencing. The ground electrode was placed superior to the nasion. We recorded the data at a 1000 Hz sampling rate.

All data was preprocessed in MATLAB using FieldTrip (Oostenveld et al., 2011). We epoched the data from –1.0 s to 3.0 seconds after stimulus onset. We applied a notch filter at 50 Hz, and all data were band-pass filtered between 0.1-100 Hz. Noisy trials were removed by visual inspection before further analysis. We performed Independent Component Analysis using the ‘runica’ implementation from EEGLAB through FieldTrip (Delorme & Makeig, 2004). Components reflecting ocular artifacts were identified from their topographies and time courses and removed. After preprocessing, participants retained an average of 159.68 out of 170 trials (SD = 16.89). An average of 2.40 ICA components were removed per participant (SD = 1.19, range = 1–6).

### Frequency tagging analyses

We performed frequency analyses separately for each participant and condition, focusing on the 0.8-1.8 stimulus window. This window was chosen because it captures the time window in which the critical verb, mouth movements, and gesture are unfolding. We calculated power spectra using fast Fourier transform and a single Hanning taper. Frequencies of interest ranged from 1 to 80 Hz in 1 Hz steps, with zero padding to 10 seconds.

We used stimulus-period local log signal-to-noise ratio (local log-SNR) as a dependent measure for later analyses. For each target frequency, local log-SNR was calculated as the log power at the target frequency minus the log mean power of selected neighboring noise frequencies. This approach quantifies a tagged response relative to neighbouring spectral bins, reducing the influence of 1/f power decay and broadband power differences (Bach & Meigen, 1999; Norcia et al. 2015). Local log-SNR was computed for 58, 63, and 65 Hz, and for the IM frequencies 2, 5, and 7 Hz. Noise bins were selected to avoid neighboring tagging or IM frequencies: 54, 55, 56, 60, and 61 Hz for 58 Hz; 59, 60, 61, 67, 68, and 69 Hz for 63 and 65 Hz; 4 and 6 Hz for 2 Hz; 3, 4, and 6 Hz for 5 Hz; and 4, 6, and 8 Hz for 7 Hz. Local log-SNR was computed at the trial level and then averaged across trials for condition-level analyses.

### Behavioral response scoring

We compared participants’ typed responses in the EEG experiment to the target sentences. Because participants typed full sentences rather than selecting from predefined alternatives, sentence-level edit similarity was used as the behavioral comprehension measure. Edit similarity was based on the Levenshtein edit distance, which quantifies the minimum number of character insertions, deletions, or substitutions needed to transform the typed response into the target sentence (Levenshtein, 1966). Before scoring, typed responses and target sentences were normalized by converting text to lowercase, removing punctuation, replacing multiple spaces with a single space. Sentence-level edit similarity was then computed as:

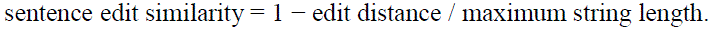

This score ranges from 0 to 1, with 1 indicating an exact match between the typed response and the target sentence. Higher scores therefore indicate better sentence-level comprehension.

### Statistical analysis

We used nonparametric cluster-based permutation statistics in FieldTrip (Maris & Oostenveld, 2007) to test for differences between the clear and degraded conditions, and conducted separate tests for each frequency of interest. For every test, the clear and degraded conditions were compared using dependent-samples t-tests at each included electrode, and the cluster-based procedure controlled for multiple comparisons across electrodes within each frequency test. Clusters were formed from electrodes exceeding the cluster threshold of p <.05. For every observed cluster, the cluster statistic was calculated as the sum of the t-values across electrodes in that cluster. To obtain a reference distribution under the null hypothesis, condition levels were randomly swapped within participants 5000 times. For each permutation, the same dependent-samples t-tests and clustering procedure were repeated, and the largest cluster-level summed t-value was kept. The observed cluster statistics were then compared to this permutation distribution to obtain Monte Carlo p-values. Clusters were considered significant at p < .05.

Exploratory informativeness analyses tested whether trial-level gesture or mouth-movement informativeness predicted the strength of the IM frequency. For each participant, condition, and IM frequency, trial-level IM strength was averaged over the six channels with the strongest average IM response for that condition and frequency. Spearman correlations were computed within participants between IM strength and gesture informativeness, and between IM strength and mouth-movement informativeness. Correlation coefficients were Fisher-z transformed and tested against zero across participants.

Finally, we tested links between behavior and brain responses by testing whether trial-level behavioral comprehension explained IM strength. Sentence-level edit similarity was correlated with trial-level IM strength at 2, 5, and 7 Hz within each participant and condition.

## Results

### Clear speech improved sentence accuracy

We first examined whether our degraded speech manipulation affected comprehension. Participants’ typed responses were quantified using sentence-level edit similarity between the typed response and the target sentence. Responses were more similar to the target in clear speech (M = .877) than degraded speech (M = .864), t(24) = –2.81, p = .010. Thus, degraded speech reliably reduced comprehension (see Figure 1)

**Figure 1:**
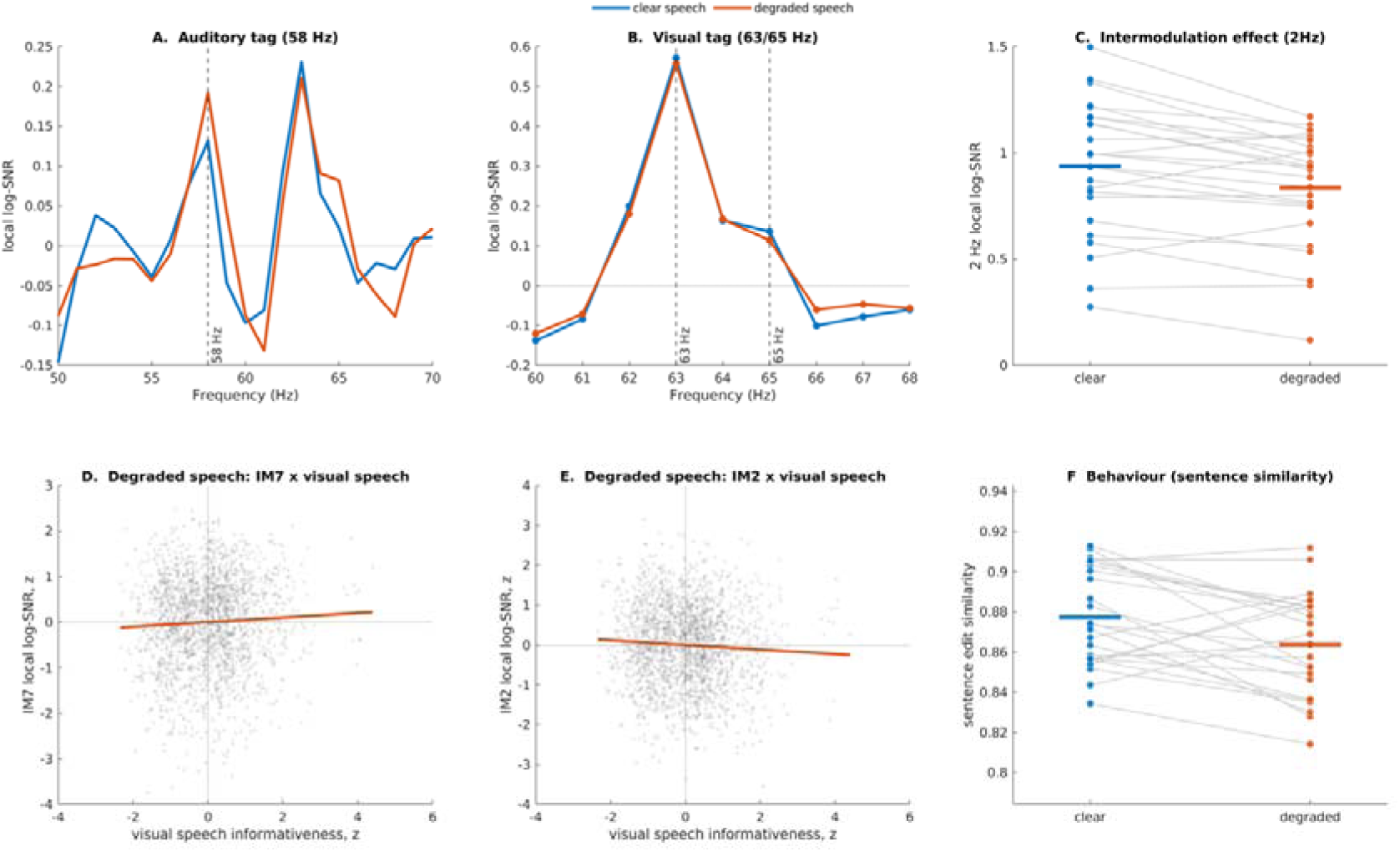
*A-B:* Grand-average local log-SNR spectra for the auditory 58 Hz tag and visual 63/65 Hz tags, shown separately for clear and degraded speech. *C*. Participant-level 2 Hz IM responses, showing stronger mouth–gesture coupling in clear than degraded speech. *D-E.* In degraded speech, visual-speech informativeness predicted IM7 positively and IM2 negatively; points show pooled within-participant z-scored trial values *F.* Sentence edit similarity was higher for clear than degraded speech.

### Degraded speech increased auditory tracking, whereas clear speech increased gesture tracking

Cluster-based permutation tests revealed a significant condition effect at the auditory tagging frequency of 58 Hz. Responses were stronger in degraded than clear speech, cluster p = .003, over a frontocentral cluster including FC1, FC2, Cz, C3, and FT9 (Figure 1). At the gesture tagging frequency of 63 Hz, responses showed the opposite pattern, with stronger responses in clear than degraded speech, cluster p = .029, over occipital channels O1 and O2 (Figure 1). No reliable condition effect was observed at the 65 Hz mouth-region tagging frequency. Thus, speech clarity modulated auditory tracking and gesture-related visual tracking, but not mouth movement tracking.

### Clear speech strengthened mouth movement-gesture coupling

We next examined whether speech clarity modulated nonlinear coupling between tagged signals. A significant condition effect was observed at the 2 Hz IM frequency, corresponding to result of the nonlinear interaction of mouth movements and gestures. Responses were stronger in clear than in degraded speech, cluster p = .027, over a cluster including CP1, P4, P8, and C4 (Figure 1). No reliable condition effect was observed at 5 Hz, indexing auditory speech-gesture coupling, or at 7 Hz (all cluster p > .05), indexing auditory speech-mouth movement coupling, demonstrating that the main IM effect of speech clarity was specific to mouth movement-gesture coupling.

### Mouth-movement informativeness, but not gesture informativeness, was related to IM strength under degraded speech

We then tested whether visual signal informativeness modulated IM strength. Gesture informativeness did not reliably predict IM responses. In contrast, exploratory analyses suggested that mouth-movement informativeness was more consistently related to IM strength, particularly during degraded speech. In degraded speech, higher mouth-movement informativeness was associated with stronger 7 Hz responses, indexing auditory speech-mouth movement coupling (p = .013). Lower mouth-movement informativeness was associated with stronger 2 Hz responses, indexing mouth movement-gesture coupling (p = .021) (see Figure 1).

### Behavioral accuracy did not robustly explain intermodulation strength

Finally, we examined whether trial-level comprehension accuracy explained IM strength. Sentence level accuracy measures did not robustly correlate with IM2, IM5, or IM7 strength (all p >.05)

## Discussion

In the present study, we investigated how listeners attend and integrate auditory speech, mouth movements, and gestures during sentence comprehension. Specifically, we asked whether the weighting and interaction of these signals are influenced by the informativeness of the visual signals and by listening conditions (clear versus degraded speech). Overall, the results revealed different patterns in neural tracking and coupling across signals. Degraded speech increased the auditory tagging response, whereas clear speech enhanced gesture-related tracking and strengthened the non-linear coupling between mouth movement and gesture. In contrast, we did not observe a reliable difference between clear and degraded speech for the mouth movement tagging frequency. Our findings suggest that multimodal comprehension is not simply characterized by stronger visual integration when speech comprehension is challenging, but that it alters the balance between auditory engagement, visual signal tracking, and multimodal coupling.

### Tracking and weighing of auditory and visual signals

Consistent with the previous RIFT studies (Drijvers et al., 2021; Seijdel et al., 2024), degraded speech elicited stronger SSEPs at the tagged auditory frequency (58 Hz) than clear speech. This finding suggests that listeners allocated more attention to the auditory signal when speech was difficult to understand. This presumably reflects increased processing demands associated with extracting linguistic information from a degraded acoustic input (Wild et al., 2012). This interpretation is further supported by the behavioral results, which showed higher sentence and verb-report accuracy in the clear compared with degraded condition, indicating that listeners were more successful at recovering the intended message when auditory input was intact, replicating previous studies (e.g., Drijvers & Özyürek, 2017).

These auditory results are in line with our behavioural results, that demonstrated that participants’ typed responses were more accurate and similar to the target sentences in clear than degraded speech. This means that the stronger response at the auditory tagging frequency might reflect increased demands on extracting linguistic information from a less reliable acoustic signal. Importantly, as we quantified tagged responses using local log-SNR, the effect reflects the extent to which the tagged auditory frequency stands out from the local spectral background, rather than it being a general broadband power increase.

In contrast, gesture-related SSEPs at 63 Hz were stronger for clear than degraded speech. This pattern of results seems counterintuitive at first, considering that attending to gestures has been shown to help disambiguate noisy speech (Drijvers & Ozyurek, 2017; Drijvers et al., 2018a, 2018b). However, this finding is consistent with previous RIFT work (Drijvers, Jensen, Spaak, 2021) that demonstrated stronger power at the tagged frequency when speech was clear. We see multiple ways to reconcile these findings. Like we put forward in previous work, we propose that since listeners do not need to extract additional information from the face during clear speech, overt visual attention might be directed to a central “resting” position on the middle of the screen when speech is clear, resulting in stronger tagging responses. Moreover, we propose that it might be useful to distinguish between the compensatory usefulness of gesture and the temporal precision with which gesture information is tracked. Gestures may be useful when speech is degraded, but degraded speech also reduced how reliable the linguistic context is in which these gestures are interpreted. In clear speech, there may be a more reliable linguistic scaffold for the listener, allowing gestures to be tracked more faithfully and consistently.

Interestingly, we did not observe a reliable difference between clear and degraded speech at the mouth movement tagging frequency. This is a remarkable finding in light of our previous eye-tracking work (Drijvers, Vaitonyte, Ozyurek, 2019), where we observed that listeners attend more to the face and mouth than to gestures when speech is degraded. However, we believe that these findings do not necessarily contradict the current findings, as overt gaze allocation and frequency-specific neural gain are related, but not the same indices of visual attention. For example, looking at the face and mouth (as was observed in our earlier eye-tracking work) might reflect strategic sampling of articulatory information, whereas the mouth movement related SSEPs at 65 Hz indices the extent to which the visual speech is neurally amplified as a signal. Our current results thus suggest that degraded speech does not produce an increase in attentional gain to mouth movements. Instead, mouth movements may have been equally attended to in both clear and degraded speech, with speech clarity affecting how representation of mouth movement was functionally coordinated with other communicative signals (see below), rather than how strongly it was tracked in isolation.

This distinction is theoretically important because attention in face-to-face communication may operate not just by amplifying individual signals, but also by regulating the coupling/relationships between signals. In a naturalistic listening context, listeners do not just attend to the mouth, the hands, or the speech separately. They must determine how information from these sources is weighted and combined as an utterance unfolds. Visual speech may therefore matter most at the level of multimodal coordination, as reflected in the IM and informativeness analyses.

### Interactions between signals

The clearest evidence for the interplay between signals was found at the IM frequency at 2 Hz, indexing interactions between the representations of mouth movements and gestures (65-63 Hz). Specifically, IM at 2 Hz responses were stronger during clear than degraded speech. This suggests that clear speech supported stronger coupling between the two visual communicative signals. This effect is consistent with ERP studies demonstrating that differences in attentional allocation can influence multisensory integration processes (Talsma & Woldorff, 2005). One possible interpretation is that clear speech provides a stable linguistic scaffold that facilitates coordinated processing of visual communicative signals. Although degraded speech may increase the need for visual information, it may simultaneously reduce the availability of attentional resources or temporal precision with which information from different modalities can be combined. In contrast, when auditory speech is reliable, listeners may be better able to align information derived from mouth movements and gestures, resulting in stronger nonlinear interactions between these visual signals.

This interpretation also suggests that informativeness of a signal and neural coupling strength are not necessarily the same. Visual signals may be more useful for comprehension when speech is degraded, but they do not necessarily show stronger coupling under degraded listening conditions. This means that RIFT IM responses are especially sensitive to temporally precise nonlinear interactions between tagged signals. Therefore, clear speech may result in stronger IM responses because the auditory signal provides a more stable contest in which visual signals can be coordinated or integrated into a coherent percept.

In contrast to IM at 2 Hz, we did not observe reliable condition effects for the IM frequencies indexing interactions between the representations of speech and gesture (5 Hz), representations of speech and mouth movements (7 Hz). The absence of robust differences between conditions does not imply that speech did not interact with mouth movement or gestures during comprehension. Rather, these interactions may be less temporally precise, more distributed over the sentence, or less detectable with the present signal-to-noise ratio. The different communicative signals may interact at distinct processing stages and timescales, making some interactions more readily detectable through IM frequencies than others.

However, interactions between speech and gesture representations were previously found when single verbs were used (Drijvers et al., 2021), suggesting that the absence of robust 5 Hz IM effects in the present study may reflect differences in task demands or stimulus complexity. When processing continuous sentences semantic information unfolds over a longer time window, allowing listeners to integrate speech and gesture information more flexibly and potentially reducing the need for temporally precise coupling between the signals. Such interactions may therefore be distributed over time. Given that IM responses are enhanced by temporal coherence between sensory signals (Xu et al., 2022), more temporally variable interactions may be less readily detectable with IM frequencies.

### The effects of gesture and mouth informativeness on the signal interactions

A key aim of the present study was to examine whether signal informativeness modulates interactions between mouth movement the communicative signals. Contrary to our expectations, gesture informativeness did not reliably predict IM strength. It is possible that the semantic similarity measures used to quantify gesture informativeness may not have fully captured the aspects of gesture meaning that listeners exploit during online language comprehension. Alternatively, gesture informativeness might only be relevant at a different or later stage of the signal integration process. Iconic gestures are thought to facilitate comprehension by aiding semantic processing (Mcneill, 1992; Morrel-Samuels & Krauss, 1992), thus gesture informativeness might affect semantically driven stages of interpretation rather than the early, temporally precise audiovisual interactions captured by RIFT. This interpretation is in line with previous finding showing that gesture speech congruence was not consistently reflected in IM responses (Drijvers et al., 2021; but see Seijdel et al., 2024).

In contrast, mouth movements are thought to facilitate comprehension by providing phonological and articulatory information that can indirectly facilitate access to semantic representations (Peelle & Sommers, 2015). As a result, mouth-movement informativeness may exert its influence at an earlier stage of the signal interaction process that is more closely aligned with the audiovisual interactions captured by IM frequencies. This interpretation is consistent with our finding that mouth-movement informativeness was more strongly associated with IM frequencies, particularly in degraded speech. Specifically, in degraded speech, more informative mouth movements were associated with stronger interactions between speech and mouth movements, suggesting that visual and auditory signals interact more strongly when the auditory signal is unreliable, and the visual speech signal is informative. Interestingly, in degraded speech, lower mouth-movement informativeness was associated with stronger interactions between mouth movements and gesture. Although speculative, this pattern may reflect a reweighting of communicative signals, whereby gesture information becomes more influential in supporting processing of mouth movements when speech is degraded and mouth-movement informativeness is reduced. Our results align with the findings demonstrating that mouth movements are especially beneficial under noisy listening conditions (Drijvers & Özyürek, 2017; Ma et al., 2009; Ross et al., 2006; Schwartz et al., 2004; Sumby & Pollack, 1954), especially when mouth movements are informative (Krason et al., 2022; Zhang et al., 2021).

More broadly, the present findings are consistent with the idea that communicative signals are weighted flexibly according to their relative informativeness and reliability, such that listeners rely more strongly on the signals that are most useful in a given context (Krason et al., 2022). Together these findings suggest that visual speech may be particularly important for supporting comprehension under adverse listening conditions, while gestures may play a compensatory role when other sources of information are less informative.

## Conclusion

Taken together, the present findings demonstrate that attentional allocation and multimodal interactions are differentially influenced by listening conditions. Degraded speech increased auditory tracking, whereas clear speech enhanced gesture processing and interactions between mouth movements and gestures. Moreover, the informativeness of mouth movements emerged as an important determinant of neural interactions when speech was noisy. These results suggest that while degraded speech increases attentional demands, clear speech appears to support stronger coordination between visual communicative signals. Taken together, our results are consistent with accounts proposing that auditory and visual information are flexibly weighted according to their relative reliability during communication (Skipper et al., 2009; Zhang et al., 2021). At the same time, our findings highlight the importance of distinguishing between the behavioral usefulness of a signal, the attentional gain applied to that signal, and the neural precision with which that signal is coordinated with other communicative signals. Our findings therefore highlight the importance of considering both signal reliability and informativeness when investigating how listeners integrate multiple sources of information during spoken language comprehension.

## Acknowledgements

We would like to thank Almut Jebens for help in collecting parts of this dataset, as well as Johan Weustink and the rest of the Technical Group at the Max Planck Institute for Psycholingistics for help setting up this experiment. This project has been supported by a Minerva Fast Track Fellowship from the Max Planck Gesellschaft, as well as the European Union (ERC Starting Grant HANDWAVE, 101219401), both granted to LD.

## Data availability statement

Data and code can be accessed via OSF: https://osf.io/u2r35/overview?view_only=45c7ddda2419468f9d0c73966c130157

